# Imaging flow cytometry enables label-free cell sorting of morphological variants from unculturable bacterial populations

**DOI:** 10.1101/2024.05.02.592280

**Authors:** Daniel Vocelle, Lindsey Thompson, Matthew P. Bernard, Nina Wale

## Abstract

Bacterial populations often display remarkable morphological heterogeneity. Flow cytometric cell sorting (often called FACS) is an important tool for understanding this diversity. FACs allows researchers to obtain pure samples of each morphological variant (or morphotype) that is present within a mixed population of cells and thus permits each morphotype to be phenotyped. In FACS, cells are first labeled with fluorescent markers such as antibodies or transgenic constructs, and then separated out based on their possession of these labels. However, since the development of fluorescent labels requires *a priori* knowledge of bacterial biology, it is often impossible to apply FACS to understudied and/or unculturable bacteria. This challenge has limited our capacity to investigate the biology of bacterial size and shape in all but a small, largely culturable subset of bacterial taxa. Here, we present an innovative strategy that permits label-free cell sorting of bacterial morphotypes, using an unculturable, pleiomorphic pathogen (*Pasteuria ramosa*) as a model bacterium. We show that imaging flow cytometry (IFC) can be used to systematically identify light-scattering and autofluorescence “signatures” of bacterial morphotypes, on which basis cell sorting can be conducted. Critically, our IFC-enabled cell sorting strategy yields samples of sufficient purity (>90%) for common downstream analyses e.g., “-omics” analyses. Our work represents an innovative application of IFC and provides an economical, widely applicable solution to a central problem in the study of bacterial diversity.

**Importance:** Bacteria come in many different shapes and sizes. Why this morphological variation exists is a longstanding question in microbiology, but it remains difficult to answer. To phenotype different morphological variants (morphotypes) within a bacterial population, we need to separate them from one another. This is normally achieved using flow cytometric cell-sorting, whereby morphotypes are labelled with fluorescent antibodies and separated on the basis of their differential fluorescence. Unfortunately, it is difficult to develop fluorescent-labels specific to unculturable or poorly studied bacteria because of the limited availability of appropriate molecular tools. Here, we demonstrate that imaging flow cytometry can be used to design and validate label-free cell sorting strategies. Recently, there has been a resurgence of interest in bacterial morphological diversity and a call to expand its study across the tree of life. Our work will help microbiologists to answer this call.

## Introduction

Bacterial populations often contain cells varying in size and shape, and this morphological heterogeneity can have a profound impact on bacterial fitness (1–3). For example, different forms of bacteria (morphotypes) may perform distinct ecological “roles” or exhibit differential susceptibility to environmental stress, permitting population persistence in the face of such stress (4–7). To fully understand the causes and consequences of morphological variation within bacterial populations, we need to extend our investigations beyond well-studied taxa and *in vitro* environments, which can suppress morphological diversity within a bacterial population (2, 8–10). Here, we present imaging flow cytometry (IFC)-enabled cell sorting, a novel procedure that allows for the purification of distinct morphotypes of unculturable bacteria living in complex milieu.

Efforts to understand the mechanistic basis and functional significance of morphological heterogeneity in a bacterial population often begin with isolating distinct bacterial morphotypes from each other and/or their environmental milieu, via flow cytometric cell sorting (or, colloquially, fluorescence activated cell sorting, FACS). However, the application of FACS to unculturable bacteria or those found in complex samples (e.g., soil, host tissues) is a challenge. A chicken-and-egg-scenario exists. In FACS, different cell populations are tagged with distinct fluorescent labels (e.g., fluorescently labelled antibodies) on whose basis the populations are separated. However, the development of morphotype-specific labels requires the characterization of morphotype-specific antigens, which itself requires that morphotypes be separated and studied. In the absence of morphotype specific labels, ‘general’ membrane permeable labels (e.g., CFSE, Mitotracker, Hoechst), which bind to molecules present in most cells (e.g., amines, DNA/RNA, or mitochondria), could be used to discriminate morphotypes in certain contexts (11). However, because general labels indiscriminately permeate all cells within a sample, they are unlikely to be helpful in distinguishing a specific morphotype if it is present in a complex, cell-rich milieu such as host-tissue or soil.

A potential solution to the problem of sorting bacterial morphotypes in the absence of labels is to exploit intrinsic variation in their light-scattering and -fluorescence properties (i.e., “light signatures”). Cells differing in size and granularity differentially scatter light, and the degree of autofluorescence may also vary among morphotypes (12). For example, viable *Bacillus anthracis* spores exhibit a distinctive autofluorescent signature upon excitation with UV light (13). Recent advances in imaging flow cytometry (IFC) simplify the identification of intrinsic light signatures of morphologically distinct cells by generating images of each cell and measures of their light-scattering and autofluorescence properties. Systematic analysis of these data allows for the identification of morphotype-specific light signatures and has been used to identify morphologically distinct subpopulations of mammalian and bacterial cells (14–17).

Here, using the pleiomorphic bacterial pathogen *Pasteuria ramosa* (hereafter, *Pasteuria*) as a model, we demonstrate that IFC can be used to efficiently design and validate strategies for label-free sorting of bacterial morphotypes*. Pasteuria* is a spore-forming pathogen of zooplankton that exhibits striking morphological variation during its life cycle (Table 1, (18)).

**Table 1.**
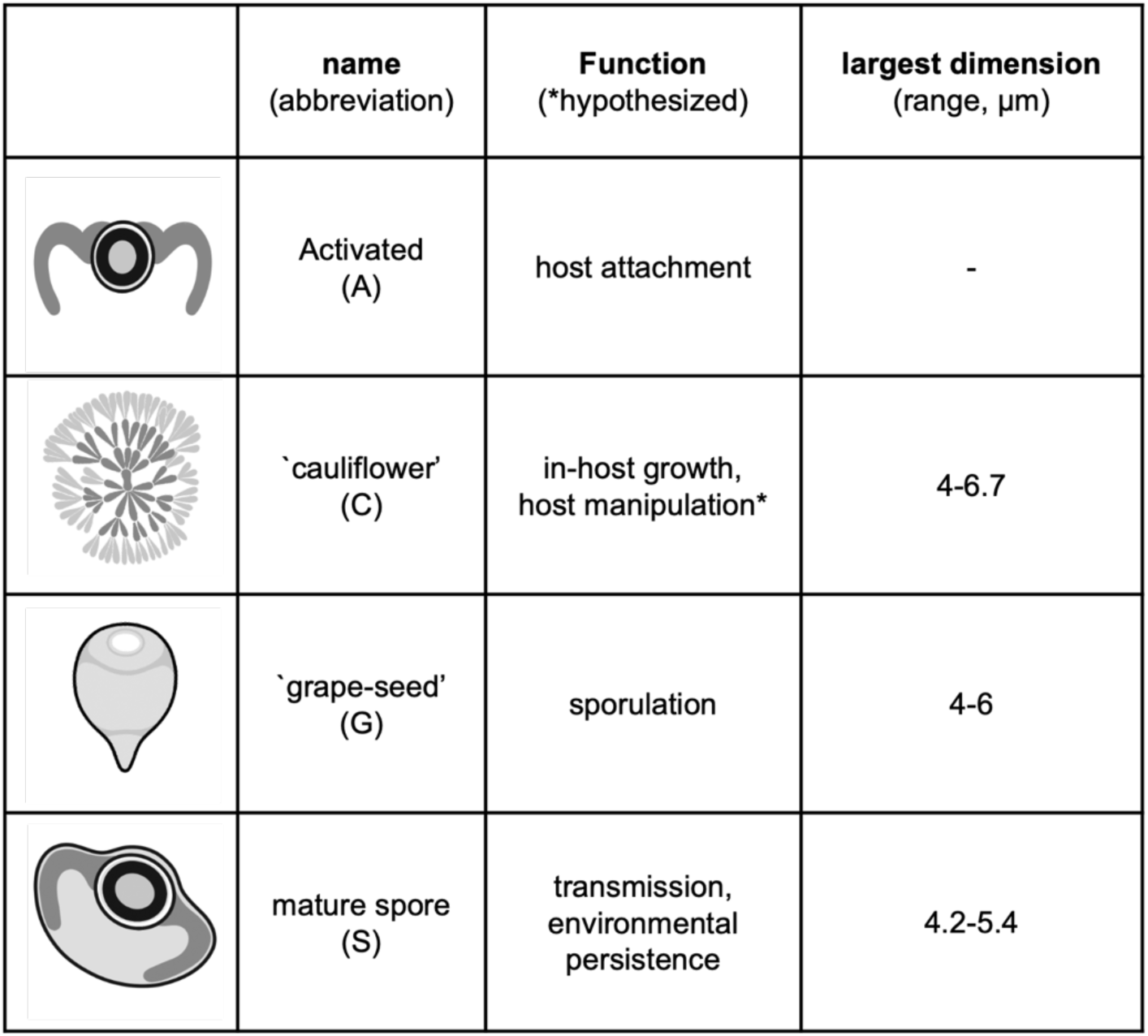
Variation in the form and function of *Pasteuria ramosa* morphotypes. Form and function information extracted from (18, 20, 21). Illustrations are schematic representations of morphotypes and are not to scale. *Indicates function is hypothesized.

| | name<br>(abbreviation) | Function<br>(*hypothesized) | largest dimension<br>(range, $\mu\text{m}$) |
| --- | --- | --- | --- |
|  | Activated<br>(A) | host attachment | - |
|  | `cauliflower'<br>(C) | in-host growth,<br>host manipulation* | 4-6.7 |
|  | `grape-seed'<br>(G) | sporulation | 4-6 |
|  | mature spore<br>(S) | transmission,<br>environmental<br>persistence | 4.2-5.4 |

Longstanding hypotheses vis-à-vis the “division of labor” among different morphotypes within the *Pasteuria* bacterial population (19) remain untested, largely because it is difficult to separate bacteria from host material for functional - omics and *in vitro* study. By combining state-of-the-art IFC instrumentation, which allows us to rapidly derive thousands of images of bacterial cells, and machine-learning algorithms, we systematically designed and validated autofluorescence-based FACS strategies that yield high-purity samples of each of four distinct morphotypes. Our work provides proof of principle that IFC-enabled sorting can be used to overcome the first barrier to understanding morphological heterogeneity in bacterial species that cannot be readily labeled.

## Materials and methods

Our goal was to design and validate cell sorting strategies suitable for the purification of 4 distinct *Pasteuria* morphotypes from the homogenate of an infected host i.e., a sample containing several *Pasteuria* morphotypes, host cells, food (algae) present in the host gut, and host microbiota. Below we describe our approach in general terms. The final procedure and results of our validation steps are described in the Results.

### Host & Parasites

Throughout, we used *Daphnia magna* (HuHO-2 genotype) hosts and *Pasteuria* (strain C1), originally collected in Hungary and Russia, respectively (22, 23). Both were generously donated by D. Ebert (U. Basel, Switzerland).

To generate samples of *Pasteuria*, we experimentally infected *Daphnia magna*. Throughout, we used a standard infection procedure. Briefly, we shook *Pasteuria* spores at 1500 rpm, overnight. On day 0 of the experiment, we filled a vessel (e.g., 24 well-plate or beaker) with media (ADaM or COMBO), added 1-10 *D. magna* <24 hours old and ∼37,500 *Pasteuria* spores per host (24). On the following day, we added a second equivalently sized dose of spores. We fed hosts increasing quantities of either *Scenedesmus obliquus* or *Ankistrodesmus falcatus* as they aged, following (25), and maintained them for 30-60 days. To diagnose *Pasteuria* infection, we assessed hosts by eye for signs of infection, including discoloration, lack of offspring and enlargement (26), and then confirmed it using microscopy.

### Instruments

We used a Thermofisher Attune CytPix flow cytometer (cytometer, hereafter) for IFC. This cytometer is equipped with a high-speed, brightfield camera (20x objective, 0.45 numerical aperture, 0.3 micron/pixel resolution), four lasers and 14 filters (Table S1). Acoustic focusing allows for the alignment of cells and particles (hereafter, events) in a narrow depth of field. As a result, this machine can acquire in-focus images an order of magnitude faster than cytometers with hydrodynamic-focusing (27).

For cell sorting, we used a BD Influx equipped with 5 lasers and 12 filters (Table S1). The default measure of fluorescence on the BD Influx is “pulse height” (rather than area). Since our goal was to use the imaging cytometer to identify fluorescence parameters that could be employed in cell sorting, we used pulse height as our measure of fluorescence on both machines.

### Data acquisition

Immediately following diagnosis, we prepared infected animals for flow cytometric analysis. First, each animal was washed in PBS (no Ca+ or Mg+). We transferred 1-4 animals, depending on the degree of infection and an initial assessment of morphotype composition via microscopy, into a 1.5mL Eppendorf tube containing PBS. We homogenized the animals briefly with an electric pestle (∼20 seconds, to avoid damaging cells) to create a slurry. We diluted the sample in 500mL Focusing Buffer (Thermofisher Scientific, catalog no. A24904) or PBS, which have equivalent refractive indices. We ran samples at 12.5mL/min until we acquired 30,000 events. Between runs, we twice flushed the cytometer line to prevent sample carryover.

### Pre-sort analysis: initial gating & image-processing

We first excluded (“gated out”) algae on the basis of their chlorophyll autofluorescence (high emission in channel R3, see Table S1). We then selected (“gated in”) the events that (i) were successfully photographed and processed and, furthermore, contained only one particle. This yielded a dataset of single events, each of which was associated with an image and a set of autofluorescence measurements.

To analyze the image data, we employed inbuilt Attune Image-Processing Software (version 6.0.1). We used the software’s bead masking model (Beads_Only_Full_Resolution_v1) which is optimized for objects 2-5mm. We chose this algorithm because (i) an alternative model optimized for objects >5mm, excluded ∼70% of our target objects and (ii) because we found that the bead model could process a variety of *P. ramosa* morphotypes, including those that exceeded 5mm (e.g., cauliflowers). Image-processing yielded a matrix containing the values of 22 imaging parameters including a variety of size and “texture” metrics (Table S2). In total, including the fluorescent parameters (i.e., pulse height, area, and width) each event was associated with 74 parameters. Before conducting further analysis, we rescaled each parameter linearly between 0 and 1.

We visualized the data using Uniform Manifold Approximation Projection for Dimension Reduction (UMAP) (28), implemented in FCS Express (De Novo Software). Briefly, this algorithm allows for the visualization of high-dimensional data in a 2D plane. By inspecting distinct regions of the UMAP, and the imaging data with which it was associated, we found that different morphotypes clustered in distinct areas of the UMAP. Preliminary analysis of this kind revealed that certain image parameters were particularly informative for distinguishing different morphotypes (Table S2). We employed this subset of parameters in further analyses, described below.

### Pre-sort analysis: identifying morphotype-specific parameters for cell sorting

For each sample, we used IFC to identify gates that would enable us to sort a specific morphotype (“target morphotype”, hereafter) from the remainder of the sample.

First, we visualized the morphological complexity of the sample using a UMAP analysis of all the scattering and fluorescent parameters, as well as several imaging parameters identified in preliminary analyses as being most-informative (Table S2, S3). We then performed cluster analysis to identify (i) a primary gate, on which basis we could distinguish our target morphotype from other bacterial cells, and (ii) subsequent gate(s), on which basis we could distinguish our target morphotype from debris (see Results). Specifically, we used FlowSOM - a neural network-like algorithm that identifies clusters of events characterized by distinct (combinations of) parameters (29). We set the consensus clustering parameter to between 8-20, depending on the sample, and otherwise used default settings (Table S3).

The above-described procedure was repeated on a sample-by-sample basis. That is, the parameters used to define the gates varied among samples (particularly in the initial stages of our work, Table S4). Due to differences in the autofluorescence of cells among samples, the exact position of the gates varied from sample to sample. Importantly, however, the gate - identification and - validation strategy - which is our focus here - remained the same across samples.

### Cell sorting

Prior to cell sorting, we transferred the sample to a 5 mL FACS tube (Corning, catalog no. 352063) and filtered it through a 100mm mesh filter. We then used the gating strategy designed using IFC (or a minor variation of the strategy, see Results, Fig. S1), to sort each of the morphotypes in the sample that were of sufficient abundance (see Table S4 for number of events sorted per sample).

### Post-sort analysis: quantification of sample purity

We verified the purity of each sorted sample using IFC. Briefly, after running the sorted sample on the imaging cytometer, the data was first processed as described above and then “down-sampled” to a subset of ∼500 events (with a few exceptions, see Table S4). Using the image data, we manually classified each of the events in the downsampled dataset as an activated-, cauliflower-, grape- or spore-morphotype of *Pasteuria* (Table 1) or as unidentified debris. Finally, we calculated the purity of the sorted sample i.e., the proportion of total events in the down-sampled dataset that were of the target morphotype. This purity measure is conservative given that we include unidentified debris such as salts in our total event count.

## Results

### IFC enables the identification of cell sorting strategies that exploit morphotype-specific light signatures

We established a robust procedure for identifying light signatures that can be harnessed for the cell sorting of morphologically distinct bacterial morphotypes, in the absence of labels. To demonstrate this approach, we provide a worked example (Fig. 1). In this example, our goal was to sort a spore (S) population from a sample comprising host (*Daphnia*) material, host-microbiota and non-target morphotypes of *Pasteuria* (cauliflower (C), grape (G)). Worked examples of the procedure as applied to activated, cauliflower and grape stages are provided in the supplement (Fig. S1).

**Figure 1:**
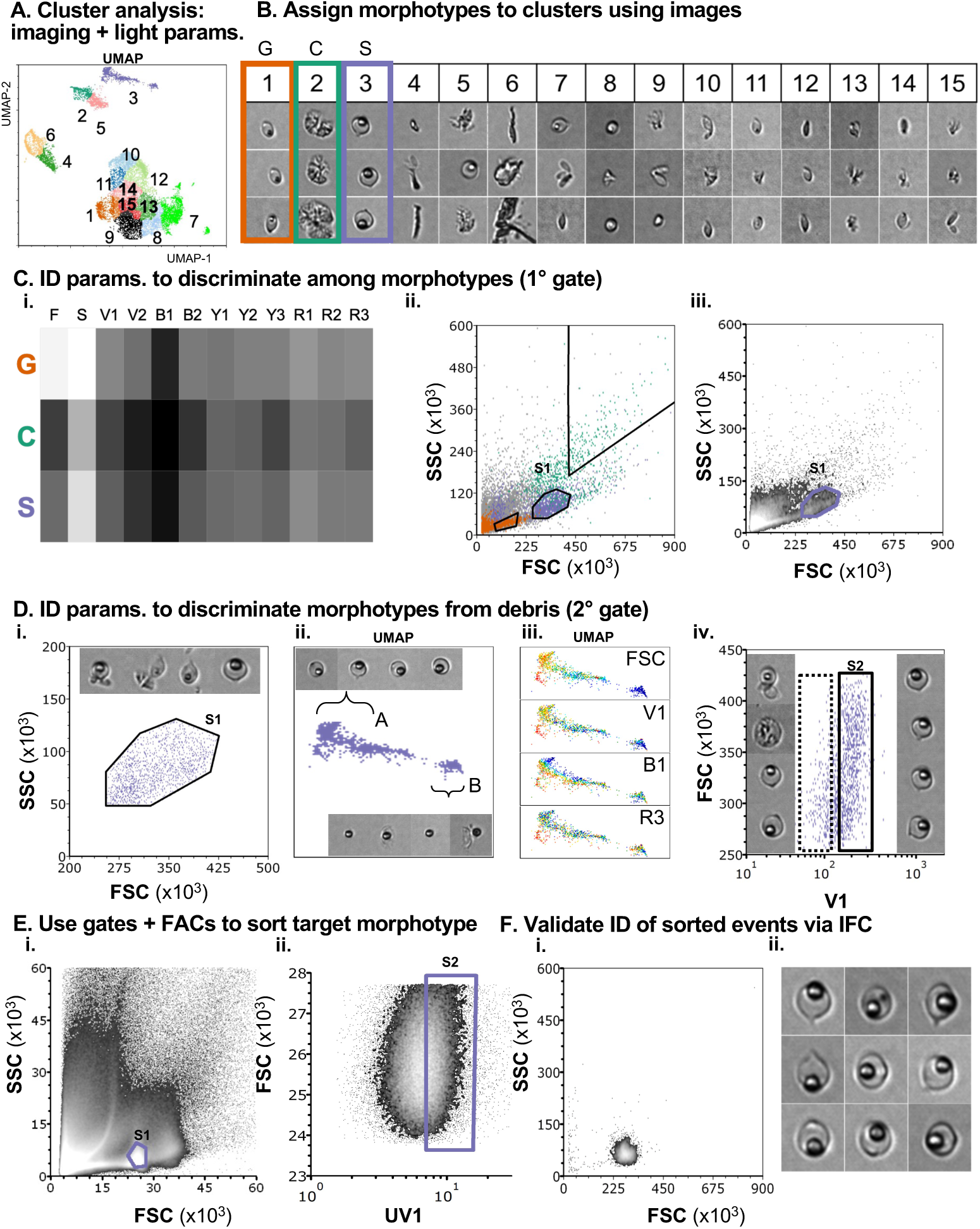
Design & validation of a label-free cell sorting strategy using imaging flow cytometry (IFC). Worked example of IFC-enabled cell-sorting, for the purification of *Pasteuria* spores (the “target morphotype”) from a sample of host material, microbiota and (non-target) *Pasteuria* morphotypes (cauliflower, C; grapes, G). **A.** Cluster analysis of imaging and light parameters (scattering, fluorescence) is performed and imposed onto a UMAP. **B.** Each cluster is assigned to a morphotype or debris using the image data. Here, clusters 1, 2, & 3 were denoted the G (orange), C (green) and S (lilac) clusters, respectively. **C.** A primary (1°) gate (S1) that distinguishes the target morphotype from non-target morphotypes is identified using (i) a “parameter heatmap”, which displays the median value of each parameter in each cluster (i; only those parameters shared among the sorter and cytometer displayed), and (ii, iii) visualization of the parameters in bipartite space (ii, events in S cluster, lilac, G cluster, orange, C cluster, green; iii, shaded from black to white to indicate increasing event density). **D.** We first inspect the images associated with the events in the 1° gate. To identify a secondary (2°) gate for separating target morphotypes from debris, regions of the target cluster that contain a pure population of the target morphotype (a) vs. a mixed population (b) are identified using the image data. Fluorescence parameters that distinguish the regions are identified by coloring the cluster by each parameter (iii, warmer colors indicate higher fluorescence intensity; note that regions A, B are differentially colored by FSC, V1 indicating the suitability of these parameters for 2° gating). Finally, events in the primary gate are plotted in bipartite space defined by the secondary parameters. The location of the secondary gate (S2) is set with respect to the image data (iv; right images, events within the gate (solid box), left images, events outside of the gate (dashed box)). **E.** The sample is analyzed using the cell sorter. It is verified that the events take approximately the same distribution on the sorter and imaging cytometer (Ei, Ciii respectively; note differences in scale); the gates are then set to the same, relative position with respect to the distribution. Here, analysis of the sample using the sorter revealed that the sorter’s UV1 channel had more discriminatory power than V1 (Fig. S3) and so we used this parameter for sorting in place of V1(Eii) **F.** To verify that sorted events are of the target morphotype and to quantify the purity of the sorted sample, the sorted sample is analyzed via IFC. First, it is confirmed that sorted events lie in expected region of bipartite space (compare Fi, Cii), then the image data is inspected and the proportion of target morphotypes quantified (Fii, representative images from sorted sample)

First, we performed a dimensionality reduction (UMAP) analysis to visualize the data in 2D space (Fig. 1A). To identify where in this 2D space each *Pasteuria* morphotype lay, we imposed the results of a cluster analysis onto the UMAP (Fig. 1A, colors/numbers indicate clusters). This cluster analysis was performed using all light-scattering and -fluorescence parameters and a subset of the imaging parameters (Table S2). We manually classified each cluster as either a morphotype-rich or debris-rich cluster using the image data (Fig. 1B). For example, since spores appeared in most of the images associated with cluster 3, we designated cluster 3 the spore-rich cluster (S-cluster; Fig. 1B, purple outline); similarly, clusters 1 and 2 were designated grape- and cauliflower-rich clusters, respectively.

Next, we used the cluster analysis to identify light-scattering and fluorescence parameters that characterized our target morphotype vs. the other morphotypes (C, G). Analysis of the median parameter values associated with each of the morphotype clusters (Fig. 1Ci), suggested that the different morphotypes could be broadly distinguished by the light scattering parameters FSC and SSC (Fig.1Ci, F,S), consistent with their differences in size and granularity (Table 1). The distribution of G-, C- and S-cluster events in FSC-SSC space confirmed this inference (Fig. 1Cii, Ciii). We thus used FSC-SSC to define our primary gate (“S1”, Fig. 1Cii).

Inspection of the images associated with the events in the primary gate revealed that it included debris as well as our target morphotype (Fig. 1Di). We therefore sought to identify a secondary gate that could be used to distinguish the target morphotype from debris. First, we inspected the images associated with the events in the target cluster, S1. This analysis revealed that there was a gradient within the cluster: on the left lay a largely homogenous population of spores (Fig. 1Dii, region A, top images); whereas on the right we found a mixed population of spores and other non-target objects (Fig. 1Dii, region B; bottom images). To identify which of the fluorescence parameters shared by the sorter and the cytometer (Table S1) could differentiate events in regions A and B we recolored the events in each cluster by their fluorescence intensity in each of the shared channels (Fig. Diii, 4 parameters shown for readability; all parameters are displayed in Fig. S2). This analysis revealed that V1 could distinguish the spore-rich region A from the mixed region B and could therefore be used in combination with FSC to define a secondary gate. Finally, we established the position of this secondary gate (“S2”) by plotting the events in our primary gate in bipartite FSC-V1 space (Fig. 1Div). We inspected the images of the events in this space (Fig. 1Div, left - images of events within dotted rectangle; right - images of events within solid rectangle) and refined the gate so that it included only events of the target morphotype (Fig. 1Div, solid rectangle).

We then performed cell sorting using the gating strategy identified via IFC. As anticipated, the sample exhibited the same qualitative, but not quantitative, pattern on the sorter as on the imaging-cytometer (Fig. 1, compare. Ei vs. Ciii). We thus positioned our gates in the same position, relative to the distribution of events, on the sorter as on the cytometer. Interestingly, we found that UV1 – a parameter available on the cell sorter but not the cytometer – could better discriminate spores from debris than V1 (Fig. S3). In this worked example, as well as other analyses where spores were the target morphotype, we used UV1 rather than V1 to define our secondary gate. With the gates set, we sorted the sample.

Finally, to validate our sorting strategy and quantify the purity of the sorted sample, we analyzed a subset of the sorted sample via IFC. First, we confirmed that the sorted events occupied the expected region of FSC-SSC space i.e., that the sorted sample occupied the same position in FSC-SSC space as the S-cluster (compare Fig. 1, Ciii, 1Fi). Second, we confirmed the identity of the sorted events using the image data (Fig. 1Fii).

### IFC-enabled sorting yields high-purity populations of each morphotype

We used the above-described gate-discovery and validation strategy to conduct label-free cell sorting on 43 occasions. These analyses involved 24 samples varying in initial composition and targeted each of the four different *Pasteuria* morphotypes.

The sorted samples ranged in purity from 75% to 99% (Fig. 2A). Of the events that were not identifiable as the target morphotype (“non-target events”), the majority were debris (e.g., salt crystals, host material, Fig. 2B). Indeed, on average the sorted sample constituted only 1.8% (range 0-8.4%) of *Pasteuria* morphotypes that were not the target (Fig. 2B).

**Figure 2:**
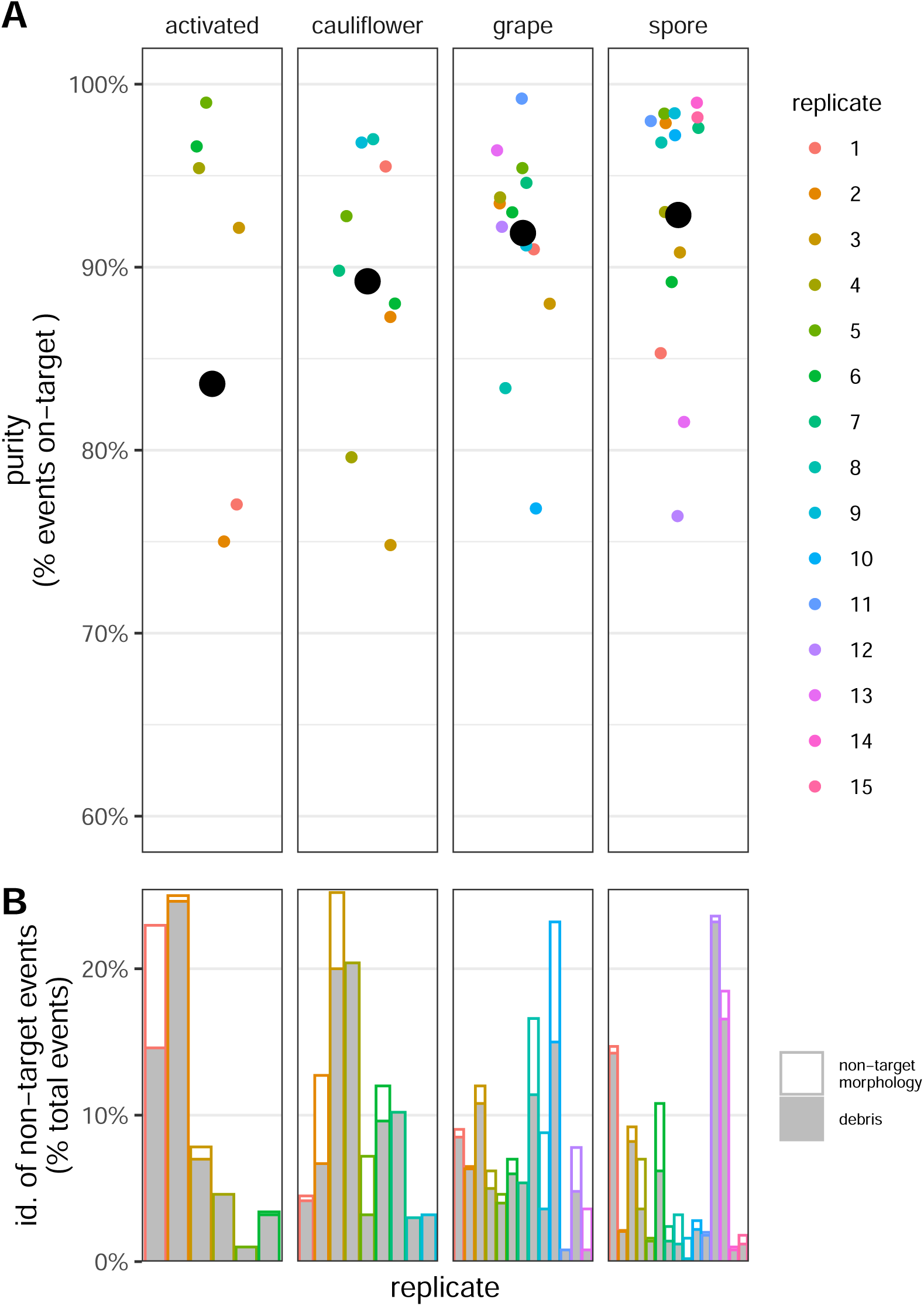
IFC-enabled cell sorting yields high purity samples of four distinct *Pasteuria* morphotypes. **A.** Purity of sorted samples, determined using image data acquired via IFC (mean purity across sorts, black, large points; purity of each replicate sort, colored, small points). Note that a replicate refers to an attempt to sort a specific morphotype i.e., replicate 1 under the “activated” and “cauliflower” columns may not have involved the same sample or been performed at the same time. **B.** Composition of the non-target fraction of the sorted sample in each replicate (colored as in **A**).

## Discussion

The first step toward understanding the physiological and ecological significance of morphological variation in a bacterial population is to obtain a pure sample of each morphotype. FACS is commonly employed for cell purification but its application to poorly studied or difficult-to-culture bacteria is limited by a paucity of morphotype-specific fluorescent-labels (antibodies, transgenics). Here, we demonstrate that IFC can be used to design and validate a label-free cell sorting strategy that efficiently separates distinct bacterial morphotypes from each other and from environmental debris. Critically, this procedure yields samples of a purity required for common downstream, “-omics” analyses i.e., >=90%. High-speed image-enabled cell sorting approaches have been innovated (30, 31) and are slowly becoming available, but their expense and limited availability makes them out of reach for most. Our methodology represents an immediately available, relatively economical alternative and could greatly expand our capacity to understand morphological heterogeneity in populations of bacteria and other organisms.

Recent innovations in the acquisition and analysis of imaging flow cytometry data are critical to the success of our approach. The first innovation is acoustic focusing. Historically, analyzing highly heterogeneous samples using imaging cytometry has been impractical, because of the slow speed of image acquisition. The acoustic focusing capability of the cytometer used herein allowed us to obtain thousands of images per second. As such, even though *Pasteuria* cells were by no means the most common events in our samples, we nonetheless obtained thousands of pictures of them. This dataset, in turn, enabled us to use data-hungry clustering algorithms to identify morphotype-specific parameters. In addition to increasing the speed with which we could acquire data, acoustic focusing also reduces the noise in the imaging and fluorescence parameters by orientating cells in a single, narrow stream (27). It is likely that this further facilitated the discrimination of morphotype-specific light signatures. Finally, acoustic focusing enabled us to rapidly validate our sorting protocol (as compared to e.g., a manual, microscopy approach) by boosting the speed with which we could inspect the morphology of our sorted events.

The second key innovation critical to our approach are machine-learning image-analysis algorithms, which are increasingly used for the analysis of high-parameter flow cytometry data (reviewed in (32, 33)). The multi-dimensionality of IFC data makes it difficult to analyze with hierarchical, manual approaches alone. Here, we used two approaches to efficiently analyze these data: the dimensionality-reduction algorithm (UMAP) allowed us to visualize the data in 2-D space, while the clustering algorithm (FlowSOM) allowed us to rapidly locate different morphotypes in this high-dimensional space and to extract their features. In future, a supervised clustering approach (e.g., (14, 34, 35)) could be used for feature-detection and may improve the efficiency of IFC-enabled gate identification.

While semi-automated computational analyses were critical to the design of our label-free cell sorting strategy, its optimization nevertheless required user-input and experience. For example, experimentation was required to elucidate the appropriate meta-clustering parameter for FlowSOM: too few clusters and the target morphotype was binned with debris/other morphotypes; too many and the target morphotype was separated into many different clusters, making it difficult to identify morphotype-specific light signatures. Similarly, user-intervention and observations were important for navigating the differences between the imaging cytometer and the cell sorter. Unlike the imaging cytometer, the cell sorter used herein does not have acoustic focusing. As a result, there was higher variability in the scattering and fluorescence data derived using the sorter vs. the imaging cytometer (compare Fig. 1Cii, Ei). We could not, therefore, position the gates in the same absolute position on the sorter as on the cytometer. Rather, we needed to manually inspect the distribution of the events and then position the gate in the same *relative* position on the sorter as on the imaging cytometer. Finally, prior knowledge vis-à-vis the UV-autofluorescence of spores (13), enabled us to better discriminate the spores from debris. These examples demonstrate that while label-free sorting is made possible by new innovations in IFC hardware and analyses, it nevertheless requires knowledge of the bacterium’s biology and the capabilities of the hardware.

Here, we applied IFC in a novel way - to purify *known* morphotypes in a bacterial population. However, a fortuitous byproduct of the IFC-enabled sorting procedure is that it yields a huge amount of image data that can be used in the more traditional application of IFC: the discovery of hitherto uncharacterized variation within a bacterial population (36). For example, in the course of our analysis, we observed a range of grape morphologies, presumably representing cells at various stages of the spore-development process, as well as events of unknown provenance. This variability is difficult to observe (let alone quantify) using microscopy, particularly when morphotypes are rare. These data may help us to address outstanding questions vis-à-vis the life cycle of this bacterial pathogen e.g., “how does *Pasteuria*’s morphology changes during the first week of infection?”.

In principle, IFC-enabled cell sorting is readily generalizable to other study systems and contexts. Nevertheless, we anticipate two hurdles that could narrow its widespread uptake. First, it may not always be possible to distinguish morphotypes on the basis of intrinsic light-scattering or autofluorescence signatures alone, particularly when hardware with limited optical components (lasers/filters) is used. Staining the sample with ‘general’, membrane permeable probes could increase the dimensionality of the data (and has proved useful in the development of a transcriptomics-based, label-free sorting approach (11)). Second, software limitations could limit the uptake of IFC-enabled cell sorting. To use IFC to identify gates requires that one can simultaneously inspect the output of machine learning analyses (e.g., cluster, UMAP analyses) and the image data. As far as we are aware, there is only one proprietary software platform (FCS express) that permits this type of analysis using data derived with the Attune CytPix. We encourage the development of open-source equivalents, particularly in languages commonly used by microbial ecologists (e.g., R, Python), to overcome this potential barrier.

Cell sorting is a first step to understanding morphological heterogeneity in bacterial populations but can only be applied to a fraction to species or contexts where that heterogeneity is exhibited. Our work offers a novel, comparatively inexpensive method that will enable microbiologists to purify – and hence phenotype – bacterial morphotypes from across the tree of life and diverse environments.

## Acknowledgements

The Attune CytPix used herein is located in the MSU Flow Cytometry Core and is supported by the Equipment Grants Program, award #2022-70410-38419, from the U.S. Department of Agriculture, National Institute of Food and Agriculture. We are grateful to members of the Wale Lab for technical assistance and to Carrie Cizauskas PhD DVM for the illustrations in Table 1.

## Contributions

DV and NW conceptualized the research. DV and LT performed the experiments.

DV designed the analysis, with consultation from MB. DV and NW analyzed the data.

NW wrote the manuscript.

DV, LT, MB edited the manuscript.

## Data availability

Description: data will be made available upon article acceptance.

