## Supplementary Material for "Imaging flow cytometry enables label-free cell sorting of morphological variants from unculturable bacterial populations"

### Supplementary Tables & Figures

| | Imaging flow cytometer (Attune CytPix) | | Cell Sorter (BD Influx) | |
|---|---|---|---|---|
| Channel | Laser | Filter | Laser | Filter |
| Ultraviolet-1 (UV1) | | | 355 | 460/50 |
| Ultraviolet-2 (UV2) | | | | 670/30 |
| Violet-1 (V1) | 405 | 450/50 | 405 | 460/50 |
| Violet-2 (V2) | | 525/50 | | 550/50 |
| Violet-3 (V3) | | 610/20 | | |
| Violet-4 (V4) | | 660/20 | | |
| Violet-5 (V5) | | 710/50 | | |
| Violet-6 (V6) | | 780/60 | | |
| Blue-1 (B1) | 488 | 530/30 | 488 | 530/40 |
| Blue-2 (B2) | | 695/40 | | 692/40 |
| Blue-3 (B3) | | | | 610/20 |
| Yellow-1 (Y1) | 561 | 585/16 | 552 | 585/29 |
| Yellow-2 (Y2) | | 620/15 | | 610/20 |
| Yellow-3 (Y3) | | 780/60 | | >750 |
| Yellow-4 (Y4) | | | | 670/30 |
| Red-1 (R1) | 637 | 670/14 | 642 | 670/30 |
| Red-2 (R2) | | 720/30 | | 720/40 |
| Red-3 (R3) | | 780/60 | | >750 |

**Table S1. Detector configuration of the imaging flow cytometer and cell-sorter used in this study**. Underlined text indicates that the two machines' parameters are identical. For each channel, the wavelength of light emitted after excitation with the laser light source and detected following separation with optical filters (center wavelength/bandwidth) is provided in nanometers. Equivalent channels are listed in the same row and given the same channel name in the main text.

| | Used in cluster analysis | Not used in cluster analysis |
|---|---|---|
| **Scatter** | Forward scatter<br>Side scatter (width, height) | |
| **Fluorescence** | Pulse "height" (all channels) | Pulse area<br>Pulse width |
| **Image**<br>*"Texture"* | Intensity (average)<br>Intensity (SD)<br>Skewness intensity<br>Kurtosis intensity<br>Entropy intensity | Normalized intensity (average)<br>Normalized intensity (SD)<br>Normalized intensity (CV)<br>Intensity (CV)<br>Max. intensity<br>Min. intensity<br>Total Intensity |
| *"Size & Shape"* | Major diameter (μm)<br>Minor diameter (μm)<br>Area ($\mu m^2$)<br>Eccentricity (%)<br>Circularity (%) | Perimeter (μm)<br>Pseudodiameter (μm)<br>Minor to major axis ratio (%) |
| *Other* | Particle count | Pixel number |

**Table S2: Parameters acquired via imaging flow cytometry and used in the cluster analysis**. Briefly, objects within images are identified via masking. The above parameters are then extracted. Intensity describes the "brightness" of each pixel of the masked object. The distribution of pixel-intensity within a particle is described by various parameters above e.g., skewness, kurtosis measures. For further details, see instrument documentation (37).

| | |
|---|---|
| UMAP | Neighbors: 50,<br>Min. Low Distance: 0.00001,<br>Iterations: 500 |
| FlowSOM | Random Cells, Linear, Gaussian, Euclidean,<br>Cluster centroids initialization method: random cells<br>Training decay function: Linear<br>2D Grid Neighborhood function: Gaussian<br>2D Grid Distance Metric: Euclidean<br>Coarse Training: 20 cycles<br>Fine-Tuning: 10 cycles, automatic neighborhood spread. |

**Table S3: Algorithm settings used in the pre-sort analysis.** Settings used represent the defaults in FCS Express analysis software package.

| | sorting | | | | | post-sort analysis | |
|---|---|---|---|---|---|---|---|
| | gate parameters (x\|y) | | | | | | |
| sample | 1° | 2° | 3° | strategy id | no. sorted events | no. events validated | purity (% on-target) |
| **spores** | | | | | | | |
| a | R3\|UV1 (1) | FSC\|SSC (1) | - | s.A (1.1.1.) | 1347 | 3000 | 85.3% |
| d | R3\|UV1 | FSC\|SSC | - | s.A | 100417 | 3000 | 97.9% |
| f | FSC\|SSC (2) | UV1\|UV2 (2) | - | s.B (2.2.1.) | 416880 | 500 | 90.8% |
| i | FSC\|SSC | R1\|UV1 (5) | - | s.C (2.5.1.) | 198207 | 500 | 93.0% |
| l | FSC\|SSC | R1\|UV1 | - | s.C | 34142 | 500 | 98.4% |
| m | FSC\|SSC | UV1\|FSC (8) | - | s.D (2.8.1.) | 29662 | 500 | 89.2% |
| n | FSC\|SSC | UV1\|FSC | - | s.D | 32657 | 500 | 97.6% |
| o | FSC\|SSC | UV1\|FSC | - | s.D | 111339 | 500 | 96.8% |
| q | FSC\|SSC | UV1\|FSC | R3\|V1 (9) | s.E (2.8.9.) | 20573 | 500 | 98.4% |
| r | FSC\|SSC | UV1\|FSC | R3\|V1 | s.E | 23099 | 500 | 97.2% |
| s | FSC\|SSC | UV1\|FSC | R3\|V1 | s.E | 85809 | 500 | 98.0% |
| t | FSC\|SSC | UV1\|FSC | R3\|V1 | s.E | 3680 | 500 | 76.4% |
| u | FSC\|SSC | UV1\|FSC | R3\|V1 | s.E | 1873 | 157 | 81.5% |
| w | FSC\|SSC | UV1\|FSC | R3\|V1 | s.E | 318508 | 500 | 99.0% |
| x | FSC\|SSC | UV1\|FSC | R3\|V1 | s.E | 50410 | 500 | 98.2% |
| **cauliflower** | | | | | | | |
| g | FSC\|R3 (3) | FSC\|UV1 (3) | R3\|FSC (2) | c.A (3.3.2.) | 5064 | 601 | 95.5% |
| n | FSC\|R3 | UV1\|SSC (9) | - | c.B (3.9.1.) | 4073 | 448 | 87.3% |
| o | FSC\|R3 | UV1\|SSC | SSC-W\|SSC (6) | c.C (3.9.6.) | 11228 | 500 | 74.8% |
| p | FSC\|R3 | UV1\|SSC | R3\|UV1 (7) | c.D (3.9.7.) | 2169 | 500 | 79.6% |
| q | FSC\|R3 | UV1\|SSC | R3\|UV1 | c.D | 1878 | 500 | 92.8% |
| r | FSC\|R3 | UV1\|SSC | R3\|UV1 | c.D | 1485 | 500 | 88.0% |
| t | FSC\|R3 | UV1\|SSC | R3\|UV1 | c.D | 6972 | 500 | 89.8% |
| u | FSC\|R3 | UV1\|SSC | R3\|UV1 | c.D | 10643 | 500 | 97.0% |
| v | FSC\|R3 | UV1\|SSC | R3\|UV1 | c.D | 10850 | 500 | 96.8% |
| **grapes** | | | | | | | |
| b | R3\|UV1 | FSC\|SSC | - | g.A (1.1.1.) | 4752 | 2457 | 91.0% |
| e | R3\|UV1 | FSC\|SSC | - | g.A | 20945 | 3000 | 93.5% |
| h | FSC\|SSC | FSC\|UV1 | SSC\|R1 (3) | g.C (2.3.3.) | 33107 | 500 | 88.0% |
| i | FSC\|SSC | UV1\|R3 (4) | FSC\|R1 (4) | g.D (2.4.4.) | 35521 | 500 | 93.8% |
| j | FSC\|SSC | UV1\|R3 | FSC\|R1 | g.D | 157471 | 500 | 95.4% |
| k | FSC\|SSC | UV1\|Y2 (6) | UV1\|R3 (5) | g.F (2.6.5.) | 16233 | 500 | 93.0% |
| l | FSC\|SSC | UV1\|Y2 | - | g.E (2.6.1.) | 5194 | 706 | 94.6% |
| m | FSC\|SSC | UV1\|B2 (7) | - | g.G (2.7.1.) | 18955 | 500 | 83.4% |
| n | FSC\|SSC | UV1\|B2 | - | g.G | 16741 | 500 | 91.2% |
| o | FSC\|SSC | UV1\|B2 | - | g.G | 213551 | 500 | 76.8% |
| q | FSC\|SSC | UV1\|SSC-W (10) | UV1\|B2 (8) | g.B (2.10.8.) | 10629 | 500 | 99.2% |
| r | FSC\|SSC | UV1\|SSC-W | UV1\|B2 | g.B | 20132 | 500 | 92.2% |
| s | FSC\|SSC | UV1\|SSC-W | UV1\|B2 | g.B | 34310 | 500 | 96.4% |
| **activated** | | | | | | | |
| c | R3\|UV1 | FSC\|SSC | - | a.A (1.1.1.) | 9191 | 3000 | 77.0% |
| f | FSC\|SSC | UV1\|UV2 | - | a.B (2.2.1.) | 2541749 | 500 | 75.0% |
| r | FSC\|R3 | UV1\|SSC-W | - | a.C (3.10.1.) | 967 | 472 | 92.2% |
| s | FSC\|R3 | UV1\|SSC-W | - | a.C | 1048 | 500 | 95.4% |
| w | FSC\|SSC | UV1\|UV2 | - | a.B | 192726 | 500 | 99.0% |
| x | FSC\|SSC | UV1\|UV2 | - | a.B | 89568 | 500 | 96.6% |

**Table S4: Gating parameters for each sort** Samples (1st column) were arbitrarily given an alphanumeric name (a-x), in ascending order of processing. Note that each sample contained 1 or more morphotypes (e.g., note that sample “n” comprised sufficient spores, cauliflowers and grapes). Each row represents an individual sort. Parameters used to define the axes (x parameter | y parameter) of the primary (1°), secondary (2°) and tertiary (3°) sorting gates are given; each unique parameter combination is given a numeric id within parentheses upon its first appearance in each column. For each target morphotype, each unique gating strategy is assigned an id (4th column); to enable comparison among sorting strategies of different morphotypes the id of the primary, secondary and tertiary gate-parameters are listed in parentheses. Of the total sorted events (no. sorted events, 6th column), a subset were imaged via IFC (no. events validated, 7th column) yielding an assessment of the purity (% on-target, 8th column) of each sort.

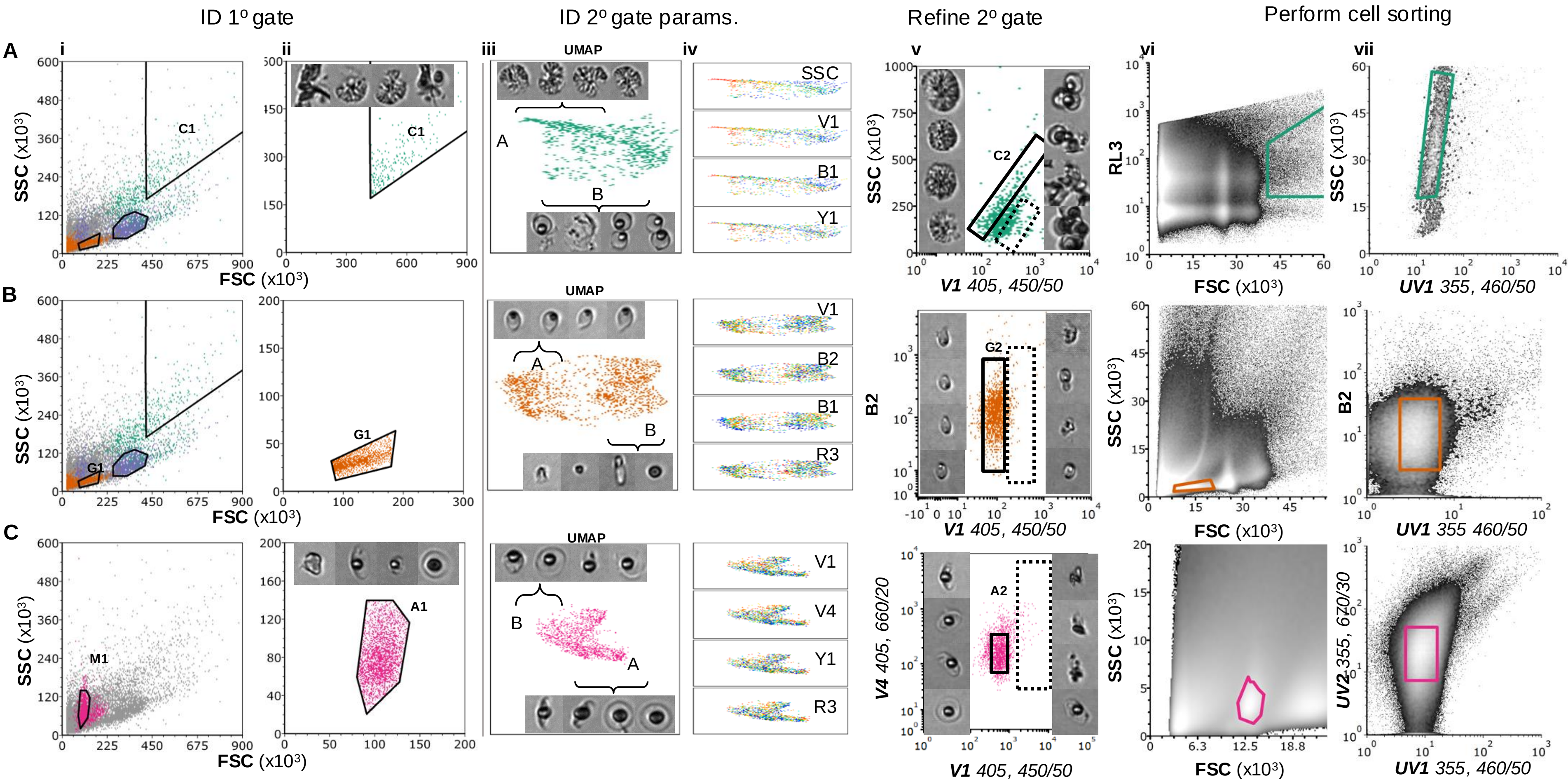


**Figure S1: Using IFC to design a strategy to sort diverse morphotypes of *Pasteuria*.** Worked example of the procedure used to cell sort cauliflower (A, green), grape (B, orange) and activated stages (C, pink) of *Pasteuria.* The analysis presented in A, B (and Fig 1) was performed using the same single sample; that presented in C used a different sample. Briefly, as described in the main text (Fig. 1A-C), we (i, ii) identified a primary (1°) gate that separated each of our target morphotypes from one another (here FSC-SSC) using the results of the cluster analysis. We then (iii) identified regions of each target cluster that contained a “pure” vs. mixed population of our target morphotype (A, B, respectively) using the image data. By (iv) coloring the clusters by each fluorescence parameter, we identified secondary parameters that could be used to discriminate target morphotypes from debris. Finally, (v) the position of the secondary gate (solid box) was set with respect to the image data (v, surrounds; images of in-gate events left, out-of-gate images right). We then performed cell sorting. Here, as in Fig. 1, the parameters used for sorting differed in a minor way from those identified via IFC (compare Ai vs. Avi; Bv vs. Bvii; Cv vs. vii); we indicate these differences in italicized text and provide laser/filter values for each of the relevant parameters. We deviated from the IFC-identified sorting strategy for one of two reasons. First, if it was possible to readily identify the target morphotype using a higher-order gate, we avoided setting additional gates. For example, here (A) we could readily distinguish cauliflower stages via their high FSC signal even using the R3-FSC gate, which is used to exclude algae in the pre-gating analysis step (see Methods). Alternatively, we amended the gating strategy when we found that events were better distinguishable using a parameter that was available on the sorter, but not on the imaging cytometer e.g., UV1 better distinguished spores, grapes and activated stages than V1 (B,C; Fig. S3).

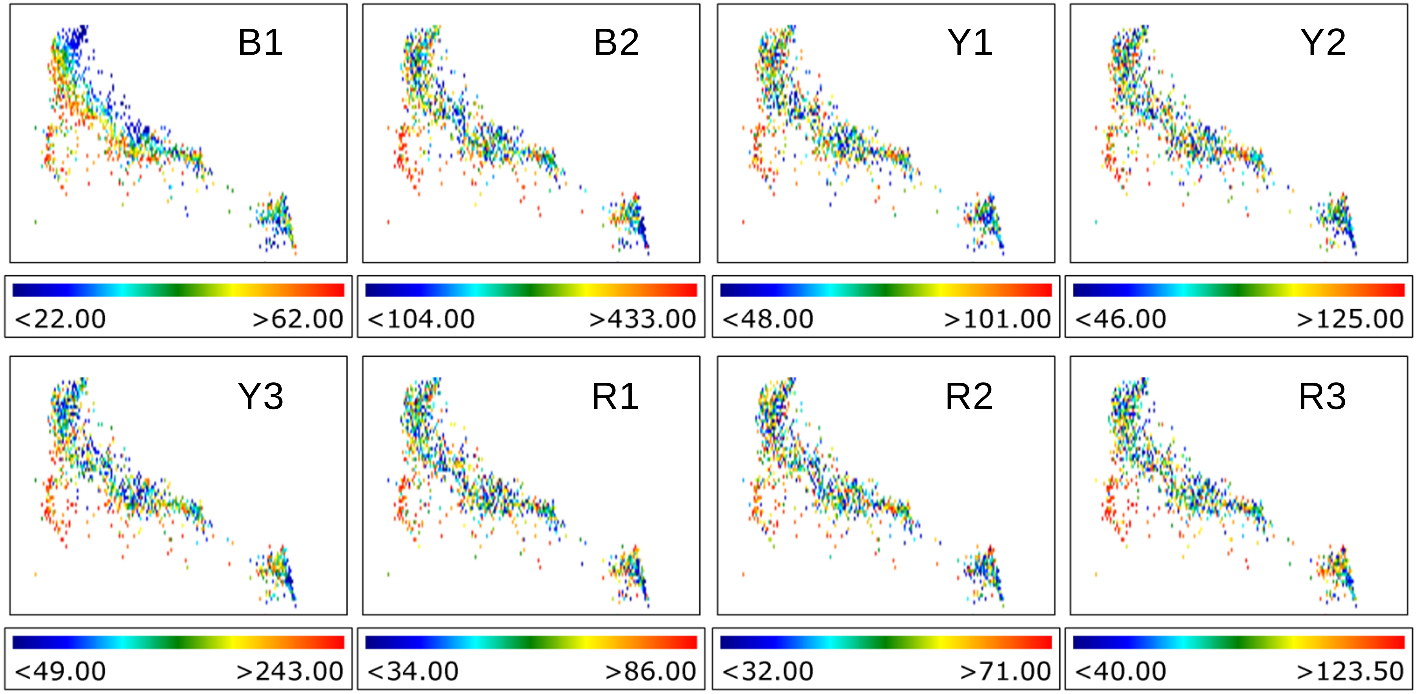


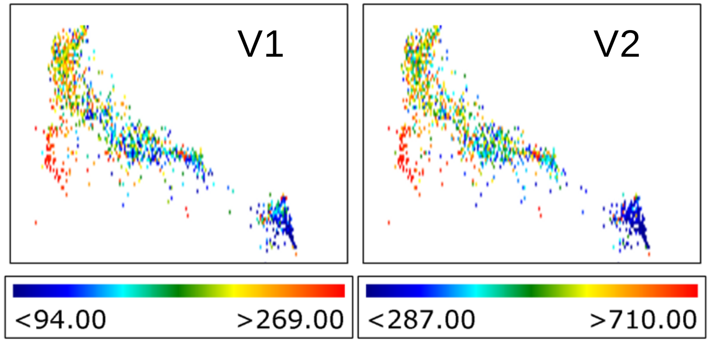


**Figure S2: UMAP analysis reveals autofluorescent signatures that can be exploited in secondary gating of *Pasteuria* spores.** Events presented in Cluster 3 in Fig. 1 of the main text, colored by the intensity of fluorescence in the 10 channels common to the sorter and cytometer (see Table S1). Each parameter was rescaled linearly so that it contained only the range of values emitted by events in the clusters; colors indicate the % intensity of each event, relative to the scale.

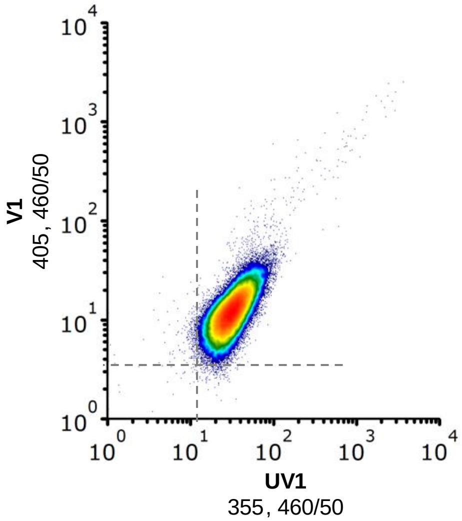


**Figure S3: *Pasteuria* spores autofluorescence with higher intensity in the UV1 than V1 channel.** Density of events in gate S1 (primary spore gate) in UV1-V1 space, as analyzed on the sorter (high density, warm colors; low density, cool colors). Dashed line imposed on the plot to enhance readability.
